# CardioShift: Resolving cell-to-cell functional heterogeneity in hiPSC-derived cardiomyocytes from high-throughput single-cell imaging

**DOI:** 10.64898/2026.10.31.763029

**Authors:** Talitha C.F. Spanjersberg, C. Nina van der Wilt, Jung-Chin Chang, Bernd Helms, Renee G.C. Maas, Joost P.G. Sluijter, Vincent Fontaine, Eric Villard, Teun P. de Boer, Rogier J.A. Veltrop, Frank G. van Steenbeek, Magdalena Harakalova

## Abstract

High-content screening of human induced pluripotent stem cell-derived cardiomyocytes (hiPSC-CMs) is widely used for disease modeling, yet standard pipelines rely on whole-well averages that obscure cell-to-cell heterogeneity. Moving from well-averages to single-cell analysis reveals hidden variations, but may introduce a pseudoreplication bias if cells are analysed as independent data points. We present CardioShift, an open-source bioimage informatics framework designed to map single-cell distributions by properly accounting for this nested variance structure. Applying this framework to 32,818 Ca^2+^ transients and 10,719 contractility-lipid profiles, we show that cell-to-cell variability represents the largest source of variance for most functional parameters, demonstrating how traditional well averages mask condition-dependent phenotypes. Furthermore, unsupervised clustering identifies condition-dependent shifts in functional subpopulations, including a distinct cellular state that accumulates lipids but maintains baseline contraction kinetics that was not detected using well- and single-cell averaging approaches. CardioShift provides a robust analytical approach to resolve hidden single-cell distributions.

## INTRODUCTION

Cells in the adult heart are not functionally uniform. Cardiomyocytes across the ventricular wall differ in electrophysiology, calcium handling and metabolism, and even neighbouring cardiomyocytes differ in size, sarcomere organisation, and protein expression^1–3^. These regional differences contribute to normal cardiac function and to disease susceptibility. Replicating this complex *in vivo* cellular environment *in vitro* remains a challenge, where human-induced pluripotent stem cell-derived cardiomyocytes (hiPSC-CMs) that are currently standard tools for disease modelling, drug screening, safety pharmacology, and genome therapy, show a similarly complex baseline heterogeneity.

HiPSC-CM-based models are typically characterised in functional multi-well assays where readouts are reported as whole-well or field-of-view averages of calcium ion (Ca^2+^) transients, contractile motion, or electrophysiological signals. These population-level measurements have proven informative and form the foundation of current standards. However, such approaches also collapse thousands of individual cell responses into a single summary metric per well, with limited insight into within-well variation. Supposedly homogeneous preparations are known to vary in size, morphology, sarcomere content, and functional behavior^4,5^, but whether and how this within-well variation shifts in response to treatment, disease genotype, or stress remains largely unexplored at scale.

To move beyond these population-level averages, high-throughput single-cell imaging has become a powerful approach^6^. However, while extracting data at single-cell resolution provides an unprecedented layer of information, evaluating these complex datasets introduces specific statistical challenges. Standard pipelines often adapt classical statistical models that treat individual cells from a shared well as independent data points. In large-scale screening settings, this generalisation can lead to pseudoreplication, inflating apparent statistical power and increasing the risk of false-positive conclusions. We reasoned that if functional heterogeneity in hiPSC-CM cultures reflects the mosaic biology of the heart, resolving it at single-cell level requires a structured, hierarchical approach. Without this resolution, effects of conditions that affect only a subset of cells, or that shift the balance between subpopulations rather than moving the entire distribution, would otherwise be underestimated or missed. Electrical coupling reduces functional differences between neighbouring cardiomyocytes *in vivo*, but whether it does so to the same extent in hiPSC-CM monolayers is unknown.

To resolve these technical and statistical bottlenecks, we developed CardioShift, an open-source bioimage informatics and analytical framework designed to decompose nested hierarchical variance structures and map single-cell functional distributions. To explore this systematically, we combined a high-throughput live-cell imaging platform with multi-parametric functional readouts in a 96-well format, measuring Ca^2+^ transients, contractile motion, and lipid content at single-cell resolution. We used lipid overload as the perturbation because it is expected to affect cells unequally.

In the adult heart, an excess supply of fatty acids contributes to lipotoxic or metabolic cardiomyopathy^7^, and individual cardiomyocytes are known to diverge in how they handle the substrate: some cardiomyocytes sequester lipids into lipid droplets, whereas others activate stress pathways that can lead to apoptosis^2,8^. Palmitate and oleate, the two most abundant circulating fatty acids, have well-documented opposing effects. Palmitate is lipotoxic at high concentrations, inducing ER stress and apoptosis through cytosolic accumulation of diacylglycerol and ceramide^9,10^. Oleate is preferentially channelled into neutral lipid storage and can even protect against palmitate toxicity^9,11^. At sub-toxic concentrations, fatty acid supplementation drives metabolic maturation, improving Ca^2+^ handling, contractile force, and mitochondrial function^12,13^. The threshold between beneficial and toxic concentrations depends on factors including cell maturation status, albumin binding, and broader medium composition^14^. At higher concentrations, particularly of palmitate, the same substrates become cytotoxic^10^. The combination of a dose-dependent maturation-to-toxicity shift and divergent effects of individual fatty acids generates a context in which both well-level and cell-level responses are expected to vary.

We applied this approach to an isogenic hiPSC-CM line pair carrying the pathogenic *MYH7* variant p.Arg403Leu (NM_000257.4(MYH7):c.1208G>T), associated with hypertrophic cardiomyopathy^15,16^ and its CRISPR-Cas9-corrected control. Recent work showed that p.Arg403Leu hiPSC-CMs are hypercontractile^17^, consistent with evidence that many HCM-associated *MYH7* variants increase force-generating myosin head availability^18,19^. Incorporating both lines allowed us to assess whether CardioShift could detect line differences alongside condition effects, and whether lipid overload interacted with the disease genotype within a nested multi-well layout.

CardioShift is a tripartite asset for the screening community as an open-source bioimage informatics pipeline to extract single-cell traces, a hierarchical statistical workflow that mathematically isolates nested variance layers to bypass pseudoreplication and a large-scale multi-modal single-cell dataset consisting of single-cell Ca²⁺ transients and single-cell contractility-lipid profiles. This resource enables the screening community to bypass statistical inflation and decode hidden subpopulation dynamics in human disease models.

## METHODS

### Preparation of fatty acid-BSA conjugates

Palmitate- and oleate-bovine serum albumin (BSA) conjugates were prepared as previously described^20^ at a 6:1 fatty acid:BSA molar ratio, corresponding to 6 mM fatty acid in 6.6% (w/v) fatty acid-free BSA. Fatty acid-free lyophilized BSA (Sigma-Aldrich, A3803) was dissolved in Milli-Q water supplemented with 1× modified balanced salt solution, and an additional 21.4 mM NaCl and 20 mM HEPES (pH 7.4). Modified balanced salt solution was generated from a sterile-filtered 10× stock containing CaCl_2_·2H_2_O (12.65 mM), KCl (45.61 mM), NaH_2_PO_4_·2H_2_O (4.49 mM), MgSO_4_·7H_2_O (4.06 mM), MgCl_2_·6H_2_O (3.94 mM), NaCl (1078.03 mM), and Na_2_HPO_4_·2H_2_O (3.36 mM).

For conjugation, 3.3 g BSA was dissolved in 25 mL Milli-Q water, 5 mL of the 10× modified balanced salt solution, 1.2 mL of 0.89 M NaCl, and 1 mL of 1 M HEPES-NaOH (pH 7.4). Palmitic acid (Sigma-Aldrich, 76119) or oleic acid (Sigma-Aldrich, O1008) was weighed into clean glass tubes at 76.93 mg or 84.74 mg, respectively, per 50 mL final conjugate. Fatty acids were saponified by adding 200 µL of 10 M NaOH, followed by 5 mL Milli-Q water preheated to 90°C, and incubating at 90°C until fully dissolved. The fatty acid-NaOH solution was then added dropwise to the BSA solution, which was maintained at 37°C while stirring on a hot plate set to 45°C. After a 10-minute incubation at 37°C to allow conjugation, the pH was adjusted to 7.4 with 1 M HCl, the volume was brought to 50 mL with Milli-Q water, and the solution was sterile-filtered through a 0.22 µm low-protein-binding PVDF filter (Millex-GV, SLGV033RB). Aliquots were stored at -20°C.

### HiPSC lines, cardiomyocyte differentiation, and culture

Human induced pluripotent stem cell (hiPSC) lines (patient-derived and isogenic control) used in this study were obtained from a patient with severe hypertrophic cardiomyopathy with diastolic dysfunction carrying a pathogenic NM_000257.4(MYH7):c.1208G>T (p.Arg403Leu) variant in the MYH7 gene as previously described^16^. Cardiomyocyte differentiation was performed using a previously published protocol^21^. After thawing, hiPSC-CMs were maintained in expansion medium containing RPMI 1640 (Thermo Fisher Scientific, 11875093), 2% Gibco B-27 Supplement (Thermo Fisher Scientific, 17504044), 2 µM CHIR (Selleckchem, S1263), and 1× penicillin/streptomycin (Gibco, 15140122)^22^. After expansion, hiPSC-CMs were replated onto Matrigel-coated clear-bottom 96-well plates (Greiner Bio-One, 655090) at 1.0 × 10^5^ cells per well to form confluent monolayers. Six replicate wells were plated per condition for *MYH7* R403L and ten replicate wells per condition for *MYH7* ISO. Recovery medium consisted of RPMI 1640 supplemented with 10% KnockOut Serum Replacement (Thermo Fisher Scientific, 10828028), 1% RevitaCell (ThermoFisher Scientific, A2644501), 2% B-27 Plus Supplement (Gibco, A3582801) and 1× penicillin/streptomycin.

### Metabolic maturation and lipid overload

Following replating, cells were allowed to recover for 24 hours and then cultured for 20 days in maturation medium^12^. The maturation medium is based on glucose-free DMEM and includes B-27 Plus Supplement (Gibco, A3582801), KnockOut Serum Replacement (ThermoFisher Scientific, 10828028), and a 1× maturation medium stock containing glucose (Sigma, G8270), L-lactate (Sigma, L7022), AlbuMAX I (albumin-bound fatty acids) (Gibco, 11020021), creatine (Sigma, C3630), taurine (Sigma, T0625), L-carnitine (Sigma, C0283), ascorbic acid (Sigma, A92902), biotin (Sigma, B4501), vitamin B12 (Sigma, V6629), non-essential amino acids (Gibco, 11140050), and penicillin/streptomycin. After the maturation period, cells were exposed to a lipid-overloaded medium for 7 days, with one medium refresh after 4 days. The control condition consisted of maturation medium supplemented with HBSS-BSA vehicle to match the BSA and buffer contribution of the fatty acid-BSA conjugates used in the lipid-overload conditions. Lipid-overload media consisted of maturation medium supplemented with 0.1 mM glycerol, 10 mM glucose, and either 600 µM palmitate, 600 µM oleate, or a 1:1 palmitate/oleate mixture (300 µM each), delivered as FA-BSA conjugates.

### Live-cell Ca^2+^ imaging

At the end of the lipid overload period, intracellular Ca^2+^ handling was visualized using Cal-520 AM (Abcam, ab171868). Cells were loaded with Cal-520 staining solution containing 0.02% Pluronic F-127 (Sigma, P2443), Cal-520 AM (Abcam, ab171868) at 1:1000 from a 2.5 mM stock, and Hoechst (Thermo Fisher Scientific, 62249) at 6 µg/ml in RPMI/B27+, after which the dye solution was diluted 1:1 with RPMI/B27 Plus Supplement before addition to the cells. Cells were incubated with 100 µL per well for 45 min, followed by the addition of Tyrode’s solution, and imaged at 37 °C in the absence of added CO_2_. Tyrode’s solution consisted of (in mM) 130 NaCl, 4 KCl, 1.8 CaCl2, 1.2 MgCl2, 18 NaHCO3, 10 HEPES and 10 glucose (pH 7.4 with NaOH).

### Live-cell WGA and BODIPY imaging

For live endpoint imaging, cells were stained in Tyrode’s solution with BODIPY 493/503 (Cayman Chemical, 25892) at 0.5 µg/ml, WGA Alexa Fluor 680 (Thermo Fisher Scientific, W32465) at 1 µg/ml, and Hoechst 33342 (Thermo Fisher Scientific, 62249) at 6 µg/ml for 15 min. The first six replicate wells were used per condition for both lines. After two washes, cells were maintained in Tyrode’s solution during imaging at 37 °C in the absence of CO_2_. WGA staining was used for membrane delineation and contraction analysis, whereas BODIPY 493/503 was used to visualize neutral lipid droplets, thereby linking lipid accumulation to functional measurements acquired from the same wells.

### Functional imaging

Single-cell functional imaging was performed on a Kinetic Image Cytometer (IC200-KIC, Vala Sciences, San Diego, USA). Cells were paced by field stimulation at 1 Hz, starting 10 s before and continuing throughout each recording. Calcium videos were recorded for 10 seconds at an acquisition frequency of 33 frames per second, and WGA/contraction videos were recorded for 5 seconds at an acquisition frequency of 70 frames per second. Calcium and contraction were measured in the same wells but not necessarily the same individual cells, as cells were not traced across sessions. For Ca^2+^ kinetics, note that we use the terms rise time and decay time rather than release and re-uptake, as the immature sarcoplasmic reticulum in hiPSC-CMs means that plasma membrane flux also contributes to the calcium transient^23^, making mechanistic attribution imprecise.

### Fixation, bleaching, and immunofluorescence

After live-cell imaging, cells were washed twice with Ca^2+^-free PBS (Gibco, 14190144), fixed for 10 min at room temperature in 4% paraformaldehyde, and then washed three times with PBS. To remove residual signal from the live-cell dyes before immunofluorescence analysis, plates were incubated for 60 minutes at room temperature under bright white light in bleaching buffer containing 3% H_2_O_2_, 20 mM NaOH, and 0.1% Triton X-100 in PBS, followed by three washes in PBS. For immunofluorescence, cells were permeabilized and blocked for 1 hour at room temperature in PBS containing 5% BSA and 0.3% Triton X-100. Primary antibodies were diluted in PBS containing 1% BSA and 0.06% Triton X-100 and incubated overnight at 4 °C. The primary antibody for ACTN2 (mouse, MilliporeSigma, A7811) was diluted at 1:200. After three washes in PBS, cells were incubated for 1 hour at room temperature with secondary antibodies diluted 1:400 in PBS containing 1% BSA and 0.06% Triton X-100. Donkey anti-mouse Alexa Fluor 568 (Thermo Fisher Scientific, A10037) was used as a secondary antibody. After immunofluorescence staining, cells were counterstained with BODIPY 493/503 at 0.5 µg/ml and Hoechst at 12 µg/ml for 10 minutes, washed, stored in PBS at 4 °C, and imaged at 20× and 40× magnification using the KIC.

### Quality control and signal classification

Cell segmentation and extraction of single-cell traces were performed in CyteSeer (Vala Sciences). Beat detection, extraction of the functional parameters, quality control and statistical analysis were performed in R using custom scripts. Calcium and contraction recordings were conducted in separate imaging sessions within the same wells. Cells were classified using two criteria applied jointly. Firstly, the normalized autocorrelation at a lag equal to the pacing period (1 s; autocorrelation at the pacing period, ACP) was computed for each trace, and the ACP distribution was thresholded using Otsu’s method. ACP is close to 1 for a cell that beats regularly at 1 Hz and low for irregular or unpaced cells. Otsu’s method places the threshold at the valley between the two peaks of the ACP distribution (0.41 for Ca^2+^ and 0.29 for contraction). This criterion implicitly rejects cells beating at frequencies other than 1 Hz, cells with aperiodic transients, and cells with too few beats to establish periodicity, since each of these reduces the autocorrelation at the 1 s lag. Secondly, a hard upper cap on detected beat count was applied: cells with more beats than expected from 1 Hz pacing over the recording duration were classified as irregular regardless of ACP (10 beats for the 10 s calcium recording, 5 beats for the 5 s contraction recording). Cells meeting both criteria were classified as electrically responsive, all others as irregular. All downstream functional analyses used only electrically responsive cells.

Two well-level gates were then applied uniformly: wells were excluded if they contained fewer than 10 electrically responsive cells, or if the fraction of cells flagged as responders by CyteSeer fell below an Otsu-derived threshold on the well-level distribution (71.0%). The second gate is needed because ACP scores rhythmicity, not signal strength. Nine calcium wells failed one or both gates, and one further well was excluded because it beat at twice the pacing frequency. For contraction, only the first gate was applied and two wells were excluded because they beat at twice the pacing frequency. Per-well values and exclusion reasons are given in **Supplementary Table 1**.

### Irregular beat quantification

The proportion of irregular cells per well was calculated for each session. Differences in the probability of irregular classification across conditions were tested using generalized linear mixed-effects models (GLMMs) with a logistic link function, fitted separately for each line (is_irregular ∼ Condition + (1 | well_uid), binomial family; lme4). The effect of condition was assessed by a likelihood-ratio test comparing the full model to a null model with only the random intercept.

### Functional subpopulation analysis

Electrically responsive cells with complete data for all clustering variables were clustered using k-means (k = 3, nstart = 50) applied to z-score standardized functional parameters. For calcium, clustering variables were: peak amplitude, Ca^2+^ transient duration at 50% and 90% recovery (CTD_50_, CTD_90_), rise time, decay time, cell area, nuclear area, and nuclear-to-cytoplasmic ratio. For contraction, variables were: contraction amplitude, rise time, decay time, nuclear area, and BODIPY intensity. Note that cell area was not available for the contraction session (CyteSeer segmented cells as circles around the nucleus in that session) and was therefore excluded from contraction clustering. Cluster stability was tested by repeating the clustering on ten random 90% subsamples of the cells. The solutions agreed closely with the full solution (adjusted Rand index ≥ 0.98 for Ca^2+^ and ≥ 0.97 for contraction). Clusters were visualized in PCA projections. Representative traces per cluster were selected as the cells closest to their cluster centroid in z-score standardized parameter space (medoid cells); mean traces across the medoid set are shown as a summary. Enrichment of clusters across conditions was tested at the well level using Kruskal-Wallis tests on the percentage of cells per cluster per well, separately for each line. The choice of k = 3 was guided by silhouette analysis and within-cluster sum-of-squares diagnostics (**Supplementary Table 2**). Silhouette width peaked at k = 2 (0.26) with a modest decrease at k = 3 (0.19). We used k = 3 because it captured a biologically informative subpopulation structure, and for consistency with the contraction analysis, where k = 3 gave the highest silhouette width (0.35).

### Statistical analysis

Functional parameters were compared across conditions using linear mixed-effects models with condition, line, and their interaction as fixed effects and well as a random intercept (param ∼ Condition × Cell_Line + (1 | well_uid)). All parameters were right-skewed (skewness > 1) and were log₁₀-transformed before model fitting. Durations longer than the 1 s pacing interval cannot be measured within one beat and were treated as missing (CTD_90_ in 157 cells and at most 12 cells for any other parameter). Residual Q-Q plots showed approximately normal residuals for all parameters (**Supplementary Fig. 1**). Fixed effects were evaluated using F tests with denominator degrees of freedom estimated by the Satterthwaite approximation. Results are reported as F(df1, df2) = F statistic, P = p-value, where df1 is the numerator degrees of freedom for the tested fixed effect and df2 is the estimated denominator degrees of freedom; therefore, df2 may be non-integer. Post-hoc pairwise comparisons were performed using estimated marginal means with Tukey correction.

To assess whether condition effects were uniform across functional subpopulations, a condition effect test was run separately within each k-means cluster. Within each cluster, a linear mixed-effects model with condition and line as fixed effects and well as a random intercept, was fitted to the cells of that cluster. Each lipid condition was compared with control using Dunnett-adjusted contrasts. Where the model could not be estimated, a Kruskal-Wallis test on well means was used instead. All analyses were performed in R (version 4.4.2) using the packages lme4, lmerTest, and emmeans.

## RESULTS

### Single-cell functional profiling of hiPSC-CMs

To test whether single-cell resolution adds information beyond conventional well-level averages in hiPSC-CM functional screening, we measured intracellular cytosolic Ca^2+^ transient and contractile motion parameters at the individual cell level in 96-well format (**Fig. 1**). As a test case, we applied a lipid overload model using an isogenic hiPSC-CM pair: one carrying a pathogenic *MYH7* variant p.Arg403Leu (NM_000257.4(MYH7):c.1208G>T; hereafter *MYH7* R403L) and its isogenic control (hereafter *MYH7* ISO). We refer to the two as hiPSC lines. Lines were generated and characterised previously, including pluripotency, genomic quality control and differentiation into cardiomyocytes^16^. In the present setup, cardiomyocyte identity was confirmed by ACTN2 immunostaining, which showed the organised sarcomeric striations characteristic of cardiomyocytes (**Fig. 2c**).

**Fig. 1.**
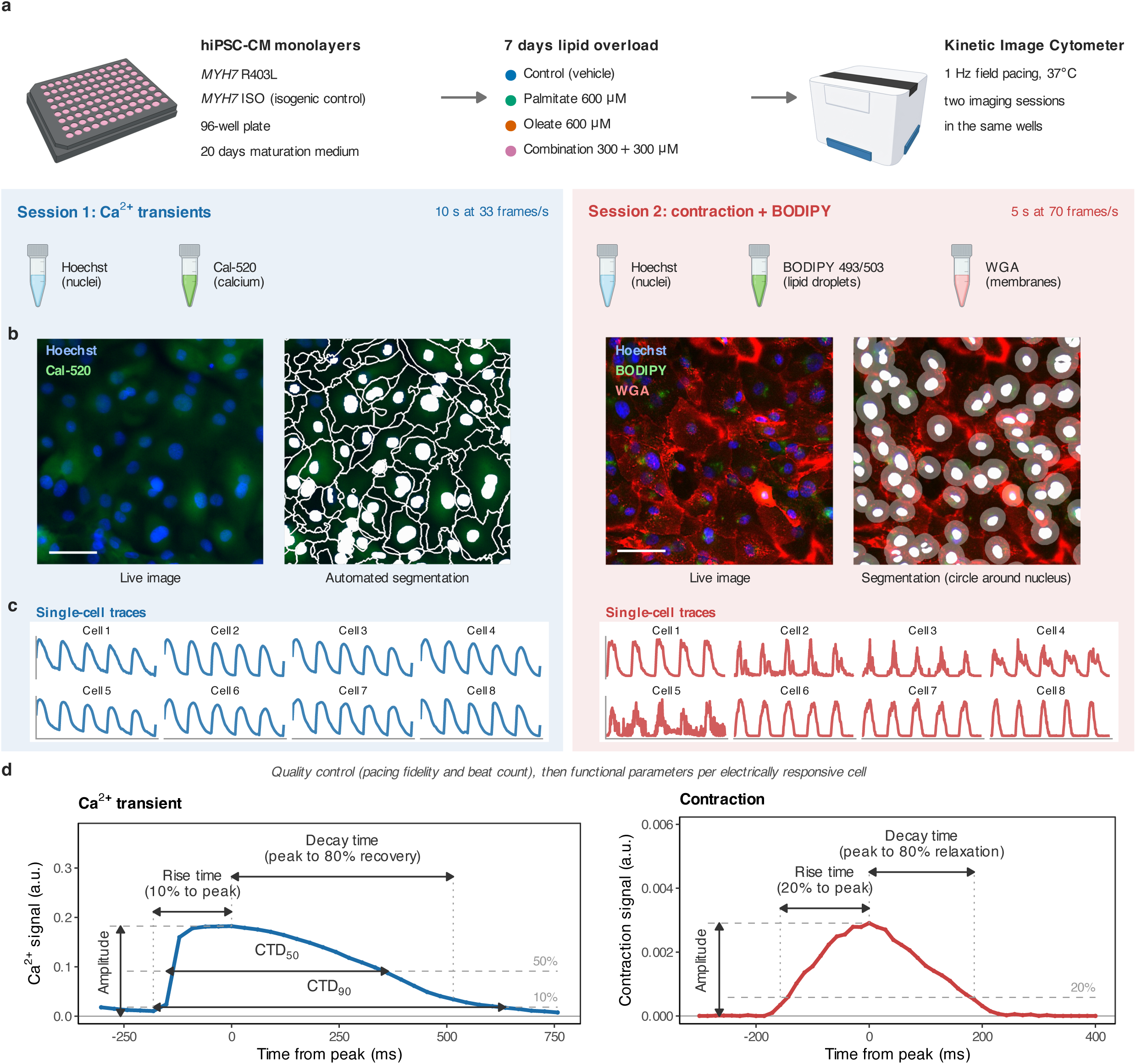
Single-cell functional imaging and analysis workflow. **a**, hiPSC-CMs carrying *MYH7* R403L and the isogenic control (MYH7 ISO) were plated as monolayers in 96-well plates, matured for 20 days in maturation medium and exposed for 7 days to control medium (BSA vehicle), palmitate, oleate, or their combination. Cells were imaged on a Kinetic Image Cytometer under 1 Hz field pacing in two imaging sessions in the same wells. **b**, In session 1, cells were stained with Hoechst and Cal-520 and Ca^2+^ transients were recorded for 10 s at 33 frames per second. In session 2, cells were stained with Hoechst, BODIPY 493/503 and WGA, and contraction was recorded for 5 s at 70 frames per second. Cells were segmented automatically from the nuclear and cytoplasmic signals in session 1 and as a circle around each nucleus in session 2. Scale bars, 50 µm. **c**, Traces of eight electrically responsive cells from one well per session (first 5 s shown). **d**, Definition of the functional parameters. Left, one Ca^2+^ transient of a control *MYH7* R403L cell, selected as the cell whose parameters were closest to the control median. Peak amplitude is the peak height of the trace. Rise time is the time from 10% of peak amplitude to the peak, and decay time the time from the peak to 20% of peak amplitude (80% recovery). CTD_50_ and CTD_90_ are the Ca^2+^ transient durations at 50% and 90% recovery, measured between the upstroke and downstroke crossings at 50% and 10% of peak amplitude. Right, contraction trace of a control MYH7 ISO cell, averaged over four beats aligned on their peaks. Rise and decay times are defined as for Ca^2+^ but at 20% of peak amplitude. Dashed lines show the reference levels as a percentage of peak amplitude. Time is shown relative to the peak. Parameters were extracted from every electrically responsive cell after quality control for pacing fidelity and beat count.

**Fig. 2.**
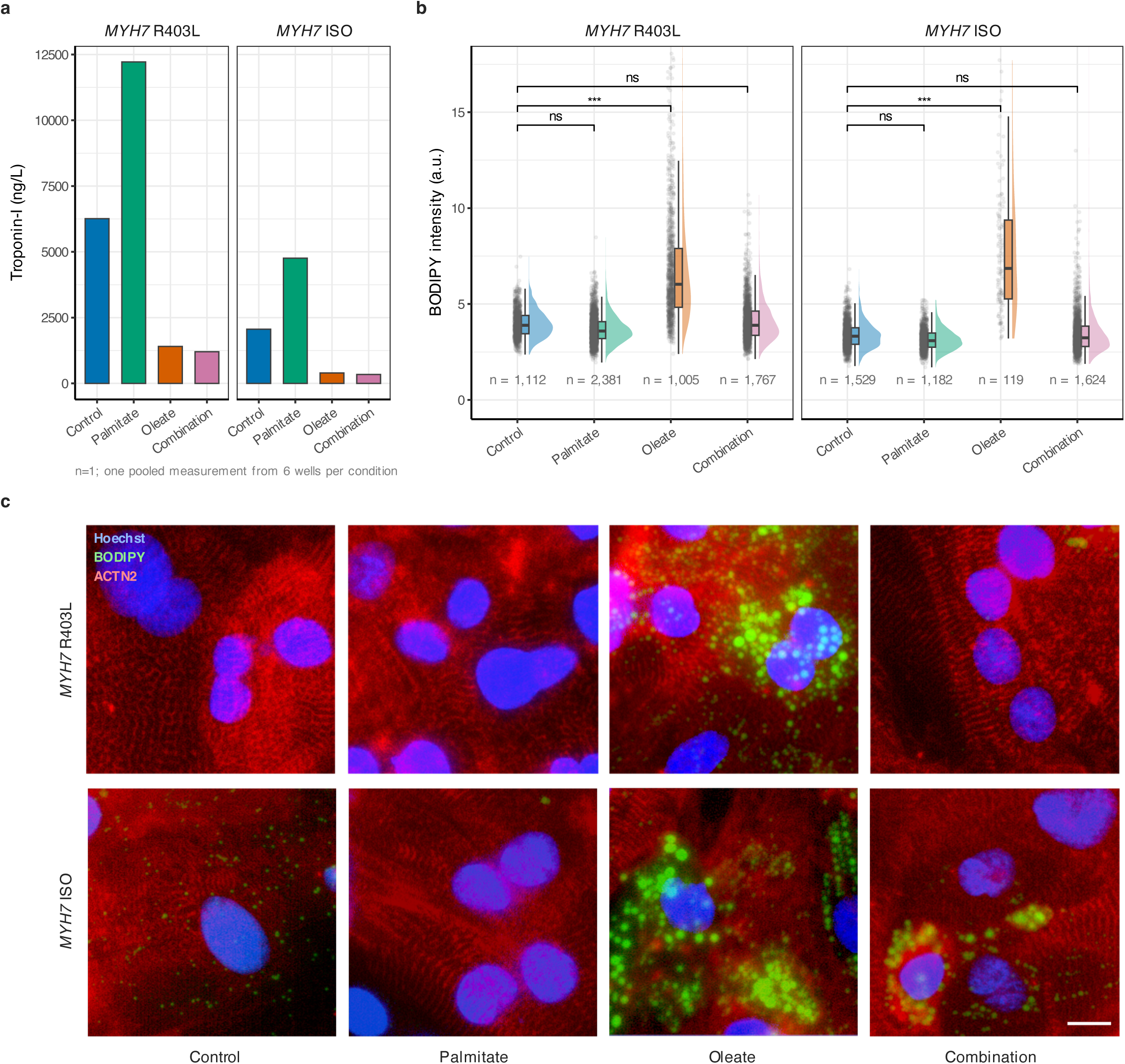
Lipid overload induces fatty acid-specific cell injury and lipid accumulation. **a**, Troponin-I release was measured from pooled supernatants of six wells per condition per cell line. Values represent single measurements per group without replicates, so no statistical test was applied. **b**, Single-cell BODIPY 493/503 intensity as a measure of intracellular neutral lipid accumulation. Distributions are shown as raincloud plots for each condition and cell line. BODIPY and contraction were measured in the same individual cells. Brackets show Tukey-adjusted post hoc contrasts against control (*** P < 0.001, ns, not significant). The y-axis shows the 0.5th to 99.5th percentile, and 54 cells above and 54 below this range are not shown. **c**, Representative immunostaining images of fixed hiPSC-CMs showing alpha actinin-2 (ACTN2, red), BODIPY 493/503 (green), and Hoechst (blue) signals. Scale bar: 10 µm.

Both lines were assayed under four conditions: maturation medium + BSA vehicle (hereafter control), 600 µM palmitate, 600 µM oleate, and a 300 + 300 µM palmitate/oleate overload combination (hereafter combination). Two imaging sessions were run on the same wells under 1 Hz field pacing, one capturing Ca^2+^ transients and one capturing contractile motion, followed by automated single-cell segmentation and trace extraction. Ca^2+^ transients and contraction were measured in the same wells but not necessarily the same individual cells, as cells were not traced across sessions. Within the contraction session, however, BODIPY lipid droplet staining and contraction were recorded in the same cells, allowing lipid content and contractile phenotype to be linked at single-cell resolution. All data derive from the same experimental run.

Across all 64 imaged calcium wells, 91.4% of segmented cells were electrically responsive, confirming that the vast majority of the imaged population consisted of beating cardiomyocytes that followed the pacing stimulus. The contraction session was markedly less robust, with 40.9% of cells electrically responsive across its 48 wells. After well-level quality control, the calcium dataset included 32,818 electrically responsive cells across 54 wells and the contraction dataset 10,719 cells across 28 wells, with 4-10 and 2-6 wells per group (**Supplementary Table 1**). Irregular beating was rare in the calcium session (0.4% of segmented cells in retained wells) and did not vary significantly across conditions in either line (GLMM likelihood-ratio test: *MYH7* R403L, chi-squared = 6.6, df = 3, P = 0.087; *MYH7* ISO, chi-squared = 7.1, df = 3, P = 0.069). In the contraction session, irregular beating was much more common (37.8% overall) and ranged from 17% (control, *MYH7* ISO) to 88% (oleate, *MYH7* ISO) (**Supplementary Fig. 2**).

Mixed-effects models with condition, line and their interaction as fixed effects and well as a random intercept, were fitted for each functional parameter. In this model, a condition effect is judged against the variation between wells, so that the hundreds of cells from one well do not count as independent replicates. Effects were tested with F tests (Satterthwaite degrees of freedom) and with Tukey-adjusted post hoc contrasts between conditions within each line. Full model output is given in **Supplementary Table 3** and all post-hoc contrasts in **Supplementary Table 4.**

### Lipid overload produces fatty acid-specific injury and lipid accumulation

Before analysing functional readouts, we confirmed that the lipid overload model produced the cellular responses expected from established biology^10–12^. Palmitate and oleate, the two most abundant circulating fatty acids, act differently on cardiomyocytes. Palmitate is broadly lipotoxic, whereas oleate is preferentially channelled into neutral lipid storage. Consistent with this, troponin-I release, measured from pooled supernatant of six wells per condition per line (one pooled measurement per group), was highest under palmitate (12,217 ng/L in *MYH7* R403L, 4,759 ng/L in *MYH7* ISO), roughly double the control baseline (6,261 and 2,061 ng/L; **Fig. 2a**). Oleate and combination conditions showed troponin-I release below baseline.

BODIPY staining, measured at single-cell level in 10,719 cells across 28 wells, showed that oleate produced by far the greatest intracellular lipid accumulation (**Fig. 2b, c**). Under oleate, mean BODIPY intensity was 1.6-fold higher than under control in *MYH7* R403L (95% CI 1.3 to 1.9) and 2.2-fold higher in *MYH7* ISO (95% CI 1.8 to 2.6), with similar differences relative to palmitate (all four contrasts P < 0.001). Palmitate and the combination were indistinguishable from control in both lines (all P > 0.7). Within oleate wells, lipid droplets were concentrated in a subset of cells (**Fig. 2c, Supplementary Fig. 3**). Both troponin-I and BODIPY data show that the four conditions produced distinct, fatty acid-specific patterns of injury and lipid loading. Palmitate caused cell damage without lipid storage, oleate caused lipid storage without damage. Having established that the model behaves as intended, we next asked what single-cell resolution adds to the functional readouts.

### Ca^2+^ transient kinetics show the strongest functional response to lipid overload

Condition effects for all parameters are summarised in **Fig. 6a** and **Supplementary Table 3**, and single-cell distributions of selected parameters are shown in **Fig. 3b**. All five Ca^2+^ transient parameters responded to lipid overload, as did nucleus area (condition main effects: F(3, 45.1 to 46.0) = 10.3 to 22.7, all P ≤ 2.5 × 10^-5^, n = 54 wells). Cell area and N/C ratio did not differ (P = 0.086 and P = 0.15). All four kinetic parameters moved in a consistent direction. Lipid loading shortened the calcium transient and slowed its upstroke. In *MYH7* R403L, decay time fell stepwise from 613 ms under control to 468 ms with palmitate, 394 ms with oleate and 341 ms with the combination, a 44% reduction at the strongest condition (P = 9.3 × 10^-7^), although the palmitate reduction alone did not reach significance (P = 0.075). Rise time moved the other way, increasing by between a third and a half (all three conditions P < 0.0001). The same responses were present in *MYH7* ISO but weaker. There, only the combination condition reached significance, reducing decay time by 24% (P = 0.008) and CTD_50_ by 23% (P = 2.1 × 10^-4^). Condition-by-line interactions for rise time (F(3, 45.9) = 10.0, P = 3.4 × 10^-5^) and decay time (F(3, 46.0) = 2.9, P = 0.046) confirm that the response was consistently stronger in *MYH7* R403L. Ca^2+^ amplitude rose under the combination condition (+35% in *MYH7* R403L, +47% in *MYH7* ISO; *MYH7* ISO control-to-combination P = 0.026, *MYH7* R403L P = 0.15), and nuclear area was smaller under every lipid condition in both lines, by 14 to 18% (significant in five of six contrasts, **Supplementary Table 4**).

**Fig. 3.**
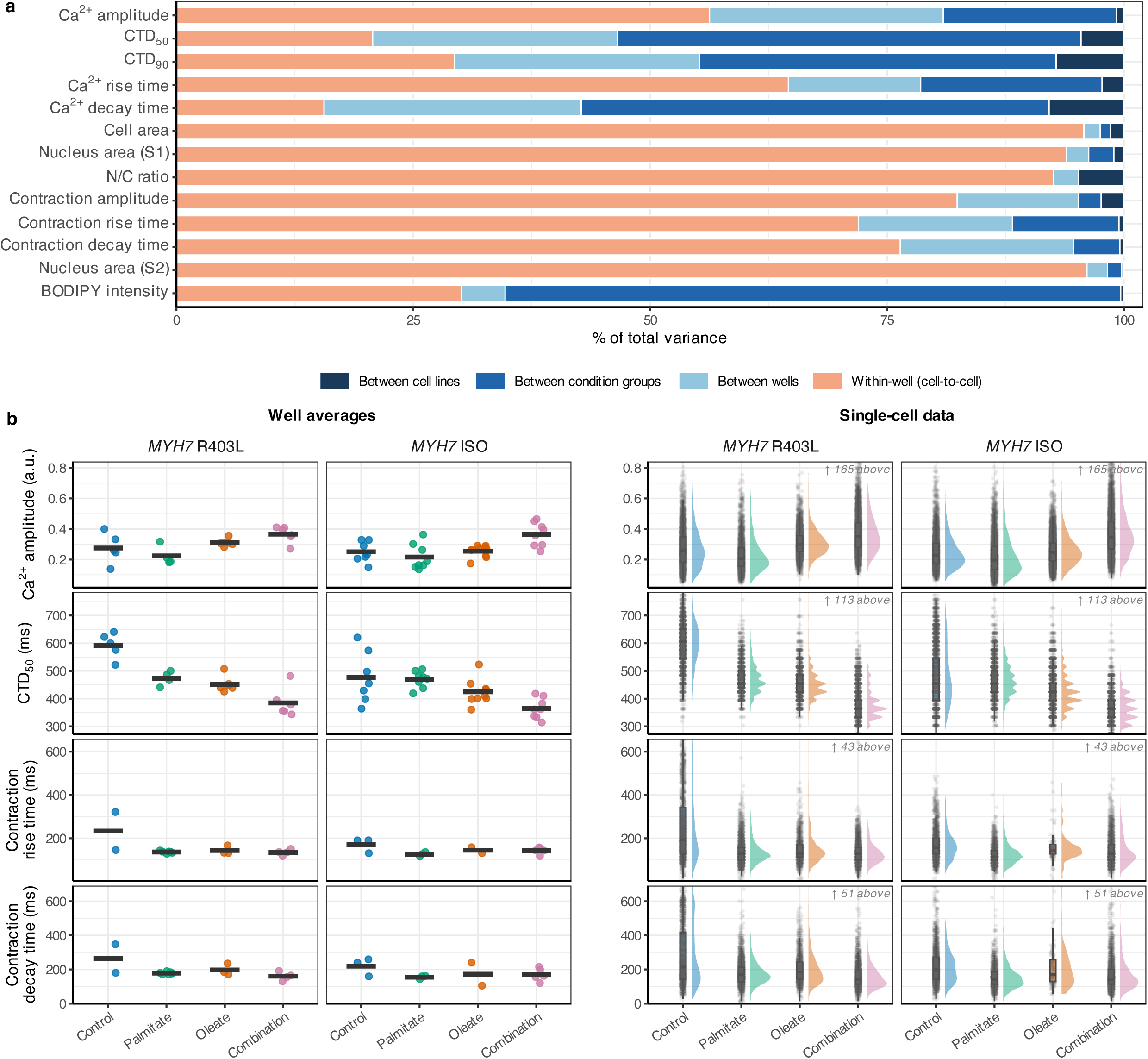
Single-cell resolution reveals within-well heterogeneity missed by well averages. **a**, Variance decomposition of all measured parameters using a linear mixed-effects model on untransformed values, with cell line as fixed effect and condition group and well as random intercepts. For each parameter, total variance is partitioned into four sources: between-cell line variance (dark blue), between-condition variance (blue), between-well variance within conditions (light blue), and within-well cell-to-cell variance (orange). **b**, Comparison of well-average (left) and single-cell (right) data distributions for four key functional parameters: Ca^2+^ transient amplitude, Ca^2+^ transient duration to 50% decay (CTD_50_), contraction rise time, and contraction decay time across two iPSC-CM lines (*MYH7* R403L, *MYH7* ISO) treated with control, palmitate, oleate, or a combination. Left panels: each point represents the mean of one well (54 calcium and 28 contraction wells); the crossbar indicates the grand mean. Right panels: violin plots show the kernel density estimate of single-cell distributions; boxes indicate median and interquartile range; individual cells shown as dots. Calcium and contraction parameters were measured in separate imaging sessions from the same wells.

Contraction parameters responded less. Contraction rise time showed a significant condition main effect (F(3, 20.7) = 4.9, P = 0.010, n = 28 wells), with post-hoc comparisons indicating shorter rise times under palmitate (P = 0.032) and combination (P = 0.038) relative to control in *MYH7* R403L. Contraction amplitude (F(3, 19.9) = 2.6, P = 0.079) and decay time (F(3, 19.7) = 3.0, P = 0.053) did not reach significance. Nucleus area in the contraction session also changed with condition (F(3, 17.1) = 4.1, P = 0.023), mainly because nuclei were smaller under oleate in *MYH7* ISO (−27%, P = 0.028). Well means of Ca^2+^ and contraction parameters were not significantly correlated (seven pairs, 23 wells, Spearman rho -0.32 to 0.39, all P ≥ 0.067, **Supplementary Fig. 5**). Independent of treatment, *MYH7* R403L had longer transient durations than *MYH7* ISO for decay time, CTD_50_ and CTD_90_ (F(1, 46.0) = 8.7 to 14.0, P = 5.0 × 10^-4^ to 0.0050) and higher mean contraction amplitude across all conditions (e.g., 0.0040 vs. 0.0029 a.u. in controls, F(1, 20.0) = 6.7, P = 0.018). *MYH7* ISO cells were larger than *MYH7* R403L cells in all four conditions (cell area 1.33-fold larger, 95% CI 1.23 to 1.43, F(1, 45.4) = 57.5, P = 1.3 × 10^-9^; **Supplementary Fig. 6**).

### Cell-to-cell variability dominates most functional parameters

Performed analyses used the well as a primary unit of observation, consistent with a well-average approach. To test how much variation this misses, we decomposed the total variance in each measured parameter into four levels. These were between lines, between conditions, between wells, and cell-to-cell within a well (**Fig. 3a**). For most parameters (9 of 13), the within-well component was largest. Cell area and nucleus area variation was almost entirely explained by the within-well range (93.9-96.1%). Contraction amplitude, rise time and decay time were also dominated by cell-to-cell variance (72.0-82.4%), with condition and line differences together contributing at most 12% to total variance. Any condition effect on these contractile parameters is therefore considered small compared with the spread between cells in the same well. Ca^2+^ parameters were more evenly distributed. Ca^2+^ decay time and CTD_50_ had the largest condition-level variance among the Ca^2+^ parameters (49.4% and 48.9%), so these are the parameters that respond most to the lipid conditions. Ca^2+^ amplitude had a substantial between-well component (24.7%), while BODIPY intensity had the highest condition-level variance (65.0%). **Fig. 3b** shows what this means in practice. Well averages leave only a few points per group and hide the shape of the distribution, which is visible in the single-cell data. Correlations between parameters within each session are shown in **Supplementary Fig. 4**.

### Unsupervised clustering reveals treatment-driven subpopulation shifts in Ca^2+^ parameter clusters

Variance decomposition shows how much cells differ but not whether they fall into distinct functional states. We therefore grouped cells with similar profiles by unsupervised clustering. K-means clustering (k = 3) was applied to z-score-standardized single-cell data for calcium (31,293 cells; 8 parameters: Ca^2+^ amplitude, CTD_50_, CTD_90_, rise time, decay time, cell area, nucleus area, and N/C ratio).

Three clusters emerged that differed mainly in transient kinetics (**Fig. 4a-d, Supplementary Table 5**). Cluster 1 (n = 13,328, 43%) had intermediate kinetics (CTD_50_ 472 ms, decay 414 ms) and the slowest upstroke (rise 228 ms). Cluster 2 (n = 13,637, 44%) had fast, high-amplitude transients (CTD_50_ 371 ms, decay 323 ms, amplitude 0.37 a.u.). Cluster 3 (n = 4,328, 14%) had slow transients (CTD_50_ 557 ms, decay 609 ms) with a fast upstroke (rise 126 ms) and larger nuclei. Cell area differed little between clusters (760 to 871 µm^2^).

**Fig. 4.**
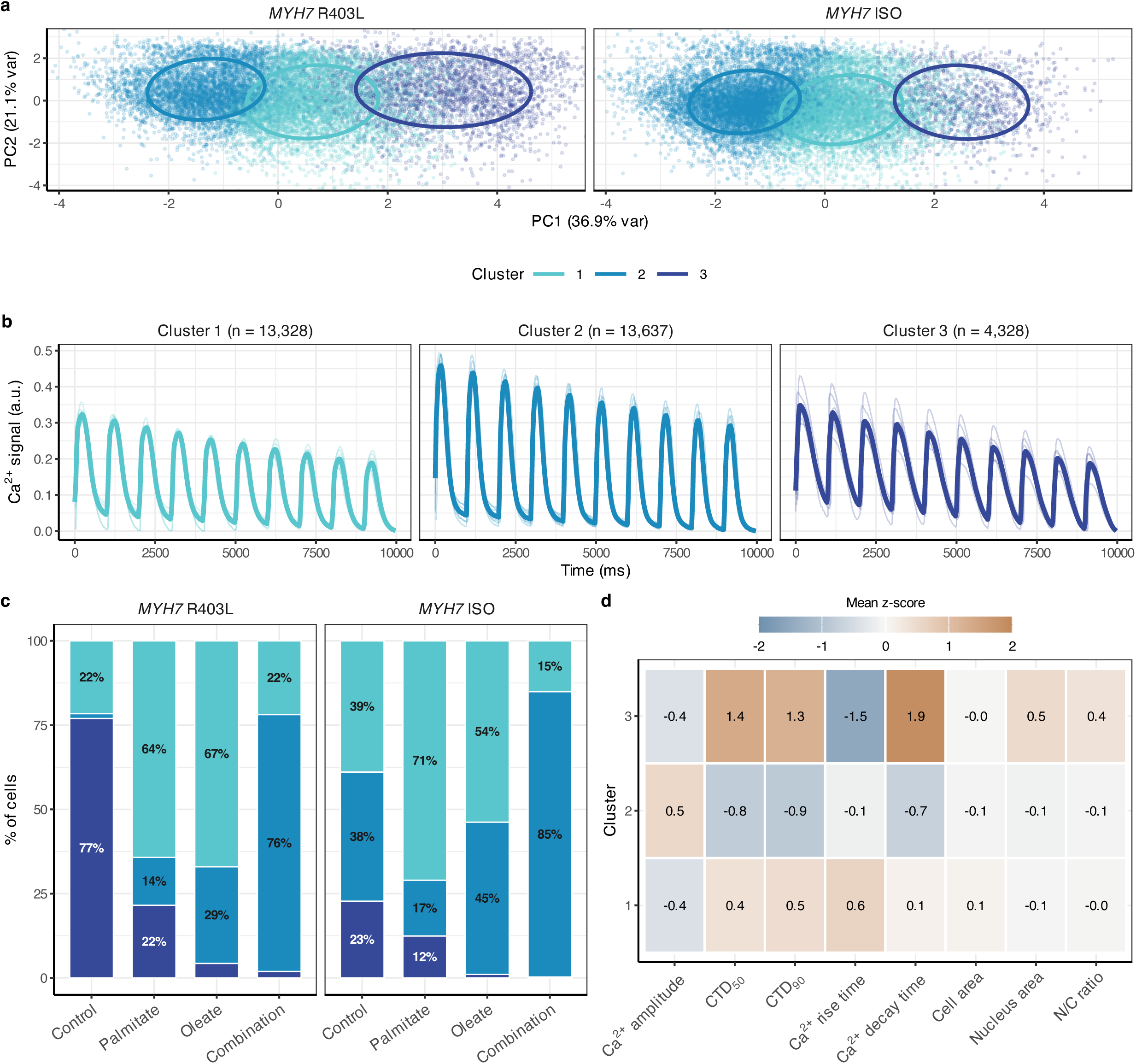
Calcium functional subpopulations shift under lipid overload. **a**, PCA projection of single-cell calcium data (31,293 cells) colored by k-means cluster assignment (k = 3). **b**, Representative Ca^2+^ transient traces for medoid cells (cells closest to each cluster centroid). Cluster 1 (light aqua): intermediate kinetics. Cluster 2 (blue): fast, high-amplitude transients. Cluster 3 (dark indigo): slow transients. **c**, Cluster composition as a fraction of total cells per condition and cell line. Composition was compared between conditions with Kruskal-Wallis tests on per-well percentages (21 *MYH7* R403L and 33 *MYH7* ISO wells; Supplementary Table 5). **d**, Cluster profiles showing the mean z-scored value of each clustering variable across clusters.

Lipid overload shifted cells between clusters (**Fig. 4c**). In *MYH7* R403L controls, 77% of cells were in the slow Cluster 3. Palmitate and oleate moved most cells to the intermediate Cluster 1 (64% and 67%), leaving 22% and 4% in Cluster 3. The combination moved most cells to the fast Cluster 2 (76%), with 2% remaining in Cluster 3. *MYH7* ISO controls were more evenly distributed (39%, 38% and 23% in Clusters 1 to 3). Here, too, palmitate and oleate enlarged Cluster 1 (71% and 54%) and the combination enlarged Cluster 2 (85%), while Cluster 3 almost disappeared under oleate and the combination (1% and 0.3%). Kruskal-Wallis tests on well-level cluster percentages showed significant condition effects for all three clusters in *MYH7* R403L (chi-squared = 10.1 to 15.0, df = 3, n = 21 wells, P = 0.0018 to 0.018) and for Clusters 1 and 2 in *MYH7* ISO (chi-squared = 17.0 and 17.8, df = 3, n = 33 wells, P = 7 × 10^-4^ and 5 × 10^-4^), but not for Cluster 3 in *MYH7* ISO (P = 0.062).

These shifts match the decay time results. Conditions that shortened mean decay time depleted the slow Cluster 3, and the combination, which shortened decay time most, enlarged the fast Cluster 2. The shift was partial, however. Under palmitate in *MYH7* R403L, mean decay time was a quarter shorter than in control, yet 22% of cells were still in the slow cluster. The mean therefore reflects a mixture of cell states rather than the phenotype of a typical cell. Resolving responses at this level can show whether a condition changes every cell or only the balance between cell states, which matters for interpreting metabolic interventions in hiPSC-CM models.

### Measuring lipid and contraction in the same cells reveals a lipid-loaded subpopulation

K-means clustering (k = 3) was applied to 10,711 cells using five parameters: contraction amplitude, rise time, decay time, nucleus area, and BODIPY intensity. Silhouette width peaked at k = 3 (0.35) and dropped at k = 4 (0.34), supporting this choice.

Three clusters emerged (**Fig. 5a-d; Supplementary Table 5**). Cluster 1 (n = 2,373, 22%) had high contraction amplitude (mean 0.0057 a.u.) and slower kinetics (rise 231 ms, decay 298 ms). Cluster 2 (n = 7,856, 73%) had low amplitude (0.0020 a.u.) and fast kinetics (rise 132 ms, decay 159 ms). Cluster 3 (n = 482, 4.5%) was defined by high BODIPY intensity (mean 9.77 a.u., compared to 3.9 and 3.7 in Clusters 1 and 2) with intermediate amplitude (0.0025 a.u.) and kinetics (rise 137 ms, decay 194 ms).

**Fig. 5.**
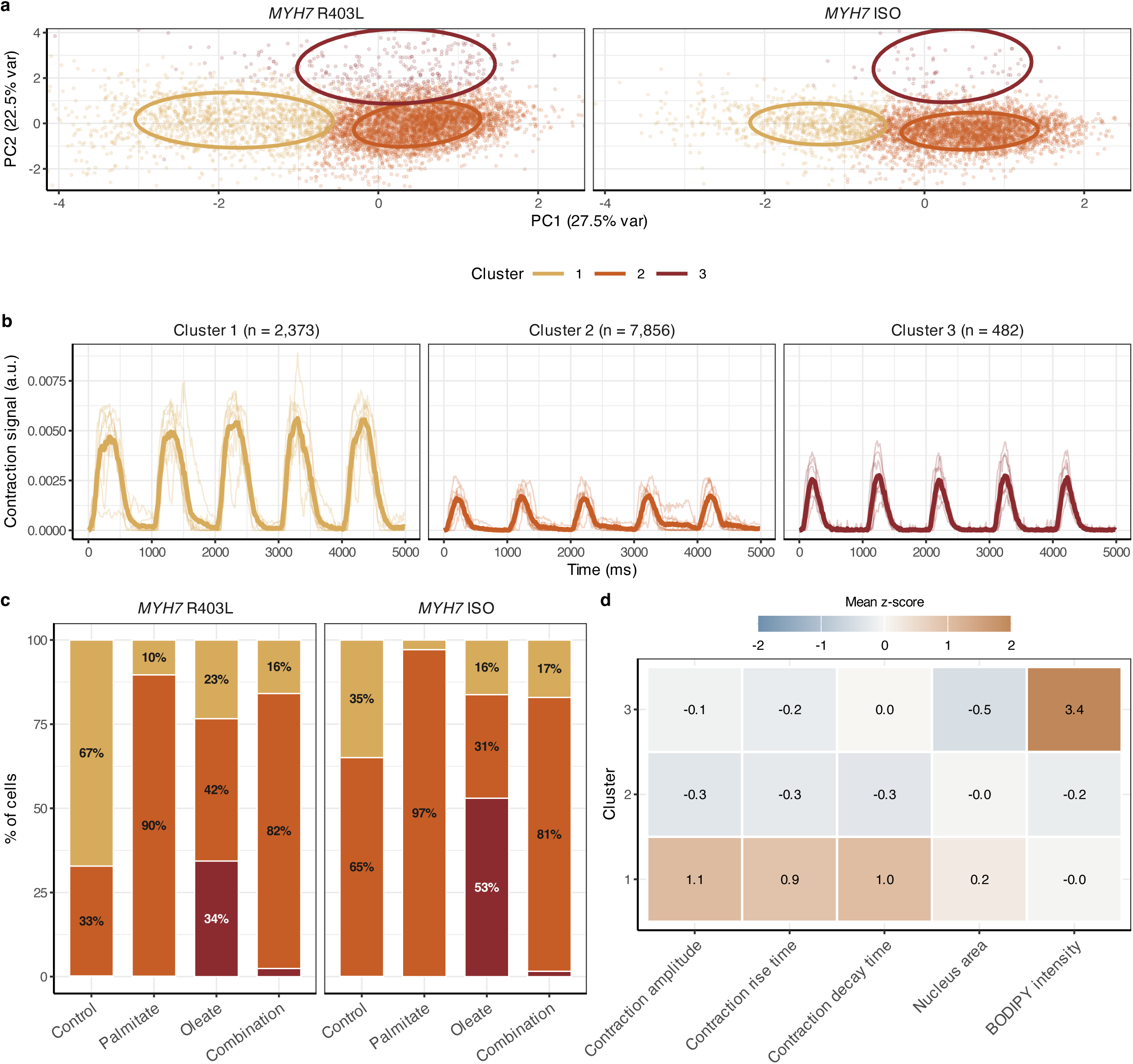
Contraction functional subpopulations and lipid-loaded cells. **a**, PCA projection of single-cell contraction data (10,711 cells) colored by k-means cluster assignment (k = 3). **b**, Representative contraction traces for medoid cells. Cluster 1 (gold): high amplitude, slow kinetics. Cluster 2 (orange red): low amplitude, fast kinetics. Cluster 3 (dark red): high BODIPY intensity with intermediate contraction profile, representing lipid-loaded cells that maintained regular beating. BODIPY and contraction were measured in the same individual cells. **c**, Cluster composition as a fraction of total cells per condition and cell line. Composition was tested as in Fig. 4 (15 *MYH7* R403L and 13 *MYH7* ISO wells). **d**, Cluster profiles showing mean z-scored values of clustering variables.

Cluster 3 emerged because BODIPY and contraction were measured in the same individual cells. These 482 cells had accumulated substantially more lipid than the broader population, yet they maintained regular contraction with a profile distinct from both the high-amplitude and low-amplitude clusters. Lower-magnification images of oleate wells show lipid droplets concentrated in a subset of cells.

Cluster composition reflected the condition. In controls, Cluster 1 (high amplitude) dominated in *MYH7* R403L (67%) but not in *MYH7* ISO (35%), consistent with the line difference in contraction amplitude. Palmitate and combination shifted cells heavily toward Cluster 2 (low amplitude, fast kinetics): 90% and 82%, respectively, in *MYH7* R403L. Oleate produced the largest shift toward Cluster 3 (high lipid content): 34% in *MYH7* R403L and 53% in *MYH7* ISO, consistent with oleate producing the greatest lipid accumulation. Every oleate well contained 15 to 53% Cluster 3 cells, against at most 0.2% in every control well. Kruskal-Wallis tests on well-level cluster percentages showed significant condition effects for Cluster 2 (chi-squared = 9.8, df = 3, P = 0.021) and Cluster 3 (chi-squared = 12.5, df = 3, P = 0.006) in *MYH7* R403L. In *MYH7* ISO, none of the individual cluster tests reached significance at P < 0.05 (Cluster 3 P = 0.063), though the limited number of wells per group (2 to 5) reduces power for this test.

### Single-cell mixed models and well-average ANOVA yield consistent results for most parameters

We compared condition main effects detected by well-average two-way ANOVA on one mean per well and single-cell mixed-effects models across all measured parameters (**Fig. 6a**). Both analyses identified significant condition effects for Ca^2+^ amplitude, CTD_50_, CTD_90_, rise time, and decay time, as well as contraction rise time, nucleus area in both sessions and BODIPY intensity. Both approaches were non-significant for N/C ratio, contraction amplitude and contraction decay time. Cell area was the only disagreement. It was significant in the well-average analysis (F(3, 46) = 10.7, P = 1.9 × 10^−5^) but not in the single-cell mixed model (P = 0.086), and no parameter was significant in the mixed model alone. Analysing every cell therefore did not inflate significance.

**Fig. 6.**
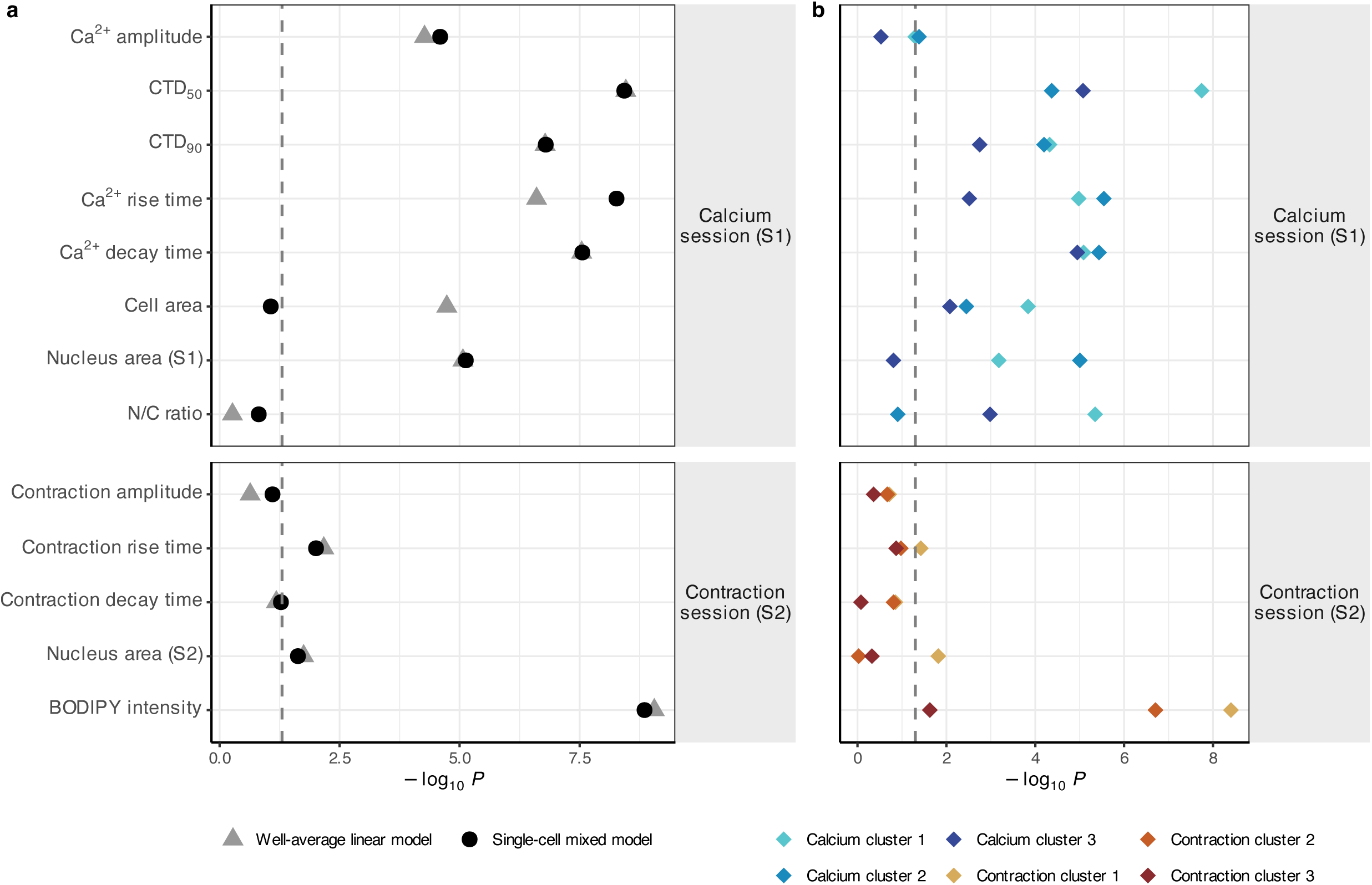
Detection of condition effects by well-average and single-cell approaches across functional parameters and subpopulations. **a**, Statistical significance (-log_10_(p)) of the lipid overload condition main effect for each measured parameter, compared between a well-average linear model (grey triangles; one mean per well, testing whether condition explains variation across wells) and a single-cell linear mixed-effects model (black circles; all individual cells included, with wells treated as a grouping factor to avoid pseudoreplication). Both models include condition, cell line and their interaction and use the same outcome scale. Dashed line: p = 0.05. Parameters are from two imaging sessions: Ca^2+^ transient parameters (amplitude; CTD_50_ and CTD_90_, the time for the Ca^2+^ signal to decay to 50% and 90% of its peak; rise time; decay time) and cell morphology (cell area; nucleus area S1, nucleus area measured in the calcium imaging session; N/C ratio, the nucleus-to-cytoplasm area ratio) from Session 1, and contraction parameters (amplitude; rise time; decay time), nucleus area (S2, measured in the contraction session), and BODIPY intensity from Session 2. **b**, The same condition effect tested separately within each k-means functional subpopulation (k = 3), using the same mixed-effects model structure as in (**a**).

When the condition effect was tested separately within each functional subpopulation (**Fig. 6b**), several parameters showed cluster-specific effects that were absent in the overall models. Cell area did not change significantly in the mixed-effects analysis but decreased within each Ca^2+^ cluster, each time under a different condition, by 29% under the combination in Cluster 1 (Dunnett P = 2.6 × 10^-4^), by 27% under palmitate in Cluster 2 (P = 0.006) and by 31% under oleate in Cluster 3 (P = 0.010). The N/C ratio, which did not change overall, rose under the combination in Cluster 1 (+38%, P = 8.8 × 10^-4^) and under oleate in Cluster 3 (+59%, P = 0.002). Because cell area and N/C ratio were themselves clustering variables, these within-cluster tests are exploratory.

## DISCUSSION

In this work, we present CardioShift as an integrated bioimage informatics and statistical resource that bridges the gap between single-cell imaging resolution and rigorous multi-well replication standards. Cardiomyocytes in the heart differ from cell to cell, and so do hiPSC-CMs in culture. However, it has been unclear how much of this variation is functional and how to analyse it without pseudoreplication. We show that single-cell analysis with a hierarchical model keeps the information that well-averages discard, while the well remains the unit of replication. The lipid overload model served as a useful test case because it produced graded, heterogeneous cellular responses rather than a uniform shift across all cells. The imaging platform used here (Kinetic Image Cytometer, Vala Sciences) was originally developed for cardiac safety pharmacology, where the emphasis was on detecting drug-induced changes in Ca^2+^ transient waveforms at single-cell level across multiwell plates^24,25^. We use the same type of data to study heterogeneity itself.

Variance decomposition showed that cell-to-cell variability within wells was the largest source of variance for most functional parameters, including contraction amplitude, rise time, decay time, cell area, and nucleus area. This framework separates variability attributable to experimental design (line, condition, well) from intrinsic cell-to-cell heterogeneity. The mixed-effects model accounts for the hierarchical data structure by allowing each well to have its own baseline, rather than assuming all wells share a common starting point, so that condition effects are tested against the variation between wells rather than between cells. By using a mixed-effect model, pseudoreplication is avoided^26^. Treating 30,000 cells from 54 wells as independent samples would make even trivial differences significant. In our data the mixed model and an ANOVA on well-means agreed for 12 of 13 parameters, and the mixed model detected no effect that the well-average analysis missed. The single disagreement, which was the cell area measurement, probably arose because the mean of a strongly skewed variable is dominated by the largest cells. The value of the mixed model is therefore not the added power, but it keeps the cell as the unit of measurement. Therefore, distributions and subpopulations can be analysed, while the well remains the unit of replication.

The clustering analysis provides a complementary view. Rather than testing for differences in means, it asks whether a condition changes the composition of functionally defined subpopulations. In the calcium data, lipid overload shifted cells away from a slow-kinetics cluster and toward a fast-kinetics cluster, consistent with the lower mean Ca^2+^ decay time results. For contraction clustering, BODIPY staining and contraction were analysed together since both measurements were obtained from the same individual cells. This co-measurement enabled the detection of a lipid-loaded subpopulation (Cluster 3) defined by high BODIPY intensity but intermediate contractile function. These 482 cells had accumulated substantially more lipid compared to the broader population, yet maintained regular contraction with a distinct kinetic profile. This cluster would be undetectable in any analysis that averages BODIPY and contraction separately across populations. It also allows lipid-loaded cells to be compared with unloaded neighbours in the same well, a comparison free of between-well variation.

As a proof-of-concept, we compared MYH7 R403L with its CRISPR-Cas9 corrected control to highlight the variant differences against its own background. The single-run configuration provides a tightly controlled, uniform hierarchical dataset that serves as an ideal baseline benchmark for evaluating CardioShift’s variance decomposition capacity. The mixed-effects model appropriately handles unbalanced designs and the total cell counts provide substantial statistical power for detecting condition effects. While subsequent independent biological validations and lab-dependent applied quality controls differ, CardioShift users may consider different thresholds for their own data interpretation.

Lipotoxic injury occurs when an oversupply of fatty acids leads to the accumulation of toxic lipid intermediates, therefore, we assessed troponin-I cellular release induced by palmitate, with troponin-I levels approximately two-fold higher than the control baseline. Oleate and combination conditions showed troponin-I levels at or below control, consistent with the established protective role of oleate in channeling fatty acids toward neutral lipid storage rather than cytotoxic intermediates^9,11^. Functional effects of lipid overload in hiPSC-CMs must be interpreted with careful attention to the metabolic context. Standard hiPSC-CM culture in RPMI/B27 provides virtually no fatty acids (only trace linoleic and linolenic acid)^27^. The maturation medium, by contrast, delivers approximately 130 µM of albumin-bound fatty acids via AlbuMAX, along with low glucose and lactate, shifting cells toward oxidative phosphorylation^12^. Using this metabolically matured baseline rather than lipid-poor RPMI/B27, means that our lipid overload conditions of 600 µM oleate or palmitate represent an excess on top of an already fatty acid-reliant metabolic state. The concentrations used here are well above the range at which fatty acids promote hiPSC-CM maturation (50 to 150 µM^12,13^) and fall within the range where palmitate-induced lipotoxicity has been documented in cardiomyocytes^9,10^.

*MYH7* R403L showed consistently higher contraction amplitude than the isogenic control (*MYH7* ISO) across all conditions. This observation aligns with the hypercontractile phenotype recently reported for R403L hiPSC-CMs using traction force microscopy^17^. Condition-by-line interactions observed for several Ca^2+^ parameters (notably rise time and decay time) suggest that the two lines respond differently to lipid overload. This is potentially relevant for understanding how genetic background modifies susceptibility to metabolic stress, but with only two lines, each cultured on its own plate, and a single experimental run, these interactions should be considered hypothesis-generating rather than definitive, especially because a high variability between lines has been reported^28,29^.

The high rate of irregular beating in the contraction session introduces additional interpretive complexity. Nearly 38% of segmented cells in the contraction session were classified as irregular, compared to less than 1% in the calcium session. Moreover, we observed a lower cell-responsiveness rate in the contraction session (40.9%) compared to the calcium handling session (91.4%). We hypothesize that this asymmetry is driven by a cumulative, condition-dependent toxicity arising from the simultaneous application of live-cell WGA membrane labeling and heavy BODIPY lipid loading. While both dyes are independently tolerated, their combination likely induces localized phototoxic stress or alters membrane mechanics, exacerbating the underlying hypercontractile phenotype of the MYH7 p.Arg403Leu mutant line. Because few or no studies have co-measured fast contraction kinetics and lipid storage simultaneously in live human cardiomyocytes, this technical trade-off represents a novel constraint of multiplexed functional-metabolic live-cell imaging. Crucially, this high-dropout scenario provided a rigorous stress-test of the CardioShift proof-of-concept, demonstrating that our hierarchical linear mixed-effects model remains stable and prevents type-I error inflation even when technical replication dropouts leave highly asymmetrical well densities.

The broader conclusion from this work is that cell-to-cell heterogeneity in hiPSC-CM populations is a highly relevant source of information. Treatments that shift subpopulation composition, affect only a fraction of cells, or interact with cell-level covariates such as size or lipid content will be missed by well-average approaches. Capturing this structure requires single-cell measurements, appropriate statistical modeling, and analytical frameworks designed to look beyond means. The workflow can be applied to other treatments, cell types and imaging readouts, because its quality control and statistical steps do not depend on the lipid model used here. Looking forward, the natural extension is to combine single-cell functional measurements with molecular readouts in the same cells. Post hoc immunofluorescence on the imaged plate is already possible, as shown here for ACTN2, but linking cells between runs requires the plate to return to the same position and the cells to stay attached. The contraction clustering in this study, in which BODIPY and contraction were measured in the same cells, illustrates what additional information is revealed when modalities converge on the same cell. Extending this principle to include Ca^2+^, electrophysiology, and molecular markers simultaneously would represent a qualitative advance over the current practice of measuring different aspects of cardiomyocyte biology in different cells and comparing them only at the population level.

Finally, by producing this multi-modal dataset, CardioShift establishes a global training and reference repository for future cardiac AI architectures and high-throughput screening tools, offering the first single-cell metabolic-functional co-measurement map to uncover masked phenotypes that serve as novel drug-discovery targets. Furthermore, the availability of paired well layouts and single-cell measurements allows computational groups to model artificial data dropouts or run down-sampling simulations to optimize statistical power guidelines for future high-content screening architectures.

## Acknowledgements

We thank Mike Hancock, Ramses Agustin, Randy Ingermanson, and Jeffrey Price at Vala Sciences for their technical support throughout this study. Their assistance with instrument setup, data acquisition, and troubleshooting was essential to the successful implementation of the Kinetic Image Cytometer in our experimental workflow.

## Author contributions

TCFS: Conceptualization, data curation, formal analysis, investigation, writing – original draft. CNW: Conceptualization, investigation, writing – review & editing. JC: Conceptualization, methodology, writing – review & editing. BH: Methodology. RM: Investigation. JS: Resources. VF: Resources. EV: Resources. TdB: Conceptualization, writing – review & editing. RJAV: Conceptualization, supervision, writing – review & editing. FvS: Conceptualization, supervision, writing – review & editing. MH: Conceptualization, supervision, writing – review & editing.

## Data availability

All scripts used to analyse the dataset are available on GitHub: https://github.com/CardiOmnics/CardioShift. Raw data tracking files are available from the corresponding author upon reasonable request.

## SUPPLEMENTARY FILES

**Supplementary Fig. 1. Residual diagnostics of the single-cell linear mixed-effects models.** Normal Q-Q plots of standardised residuals from the linear mixed-effects models (parameter ∼ Condition × Cell_Line + (1 | well_uid)) for each single-cell parameter. **a-h**, Calcium imaging session (session 1): Ca^2+^ amplitude (**a**), CTD_50_ (**b**), CTD_90_ (**c**), rise time (**d**), decay time (**e**), cell area (**f**), nucleus area (**g**) and nucleus-to-cytoplasm (N/C) area ratio (**h**). **i-m**, Contraction imaging session (session 2): contraction amplitude (**i**), contraction rise time (**j**), contraction decay time (**k**), nucleus area (**l**) and BODIPY intensity (**m**). Red lines indicate the expected distribution under normality. All parameters were modelled on the log10 scale. Each panel states the skewness of the residuals and the number of cells (n).

**Supplementary Fig. 2. Irregular cells per well in calcium and contraction sessions.** Each dot represents one well. Red triangles indicate wells excluded from the main analysis (failed well-level QC). Irregular cell percentage was generally below 2% in the calcium session across all conditions and cell lines. In the contraction session, irregular percentages were substantially higher and more variable, with the oleate condition in the isogenic control line (*MYH7* ISO) showing the highest rates.

**Supplementary Fig. 3. Lipid overload with oleate induces lipid accumulation.** Representative live-cell images of hiPSC-CMs showing BODIPY 493/503 (green) and Hoechst (blue) captured during contraction measurements. Scale bar: 50 µm.

**Supplementary Fig. 4. Spearman correlation heatmaps for Ca^2+^ and contraction parameters at the single-cell level.** Colors indicate Spearman’s rank correlation coefficient (ρ; -1 to 1); blue = negative correlation, white = none, orange = positive. Only electrically responsive cells with complete data are included, and values are shown where P < 0.05. **a**, Calcium session (S1), Ca^2+^ transient and morphology parameters. **b**, Contraction session (S2), contraction parameters, nucleus area and BODIPY intensity. CTD_50_/CTD_90_, Ca^2+^ transient duration to 50%/90% recovery; BODIPY, mean BODIPY fluorescence intensity.

**Supplementary Fig. 5. Well-level cross-modal correlations between calcium and contraction parameters.** Spearman correlations across 23 wells with paired calcium and contraction data. Seven parameter pairs were tested and none reached significance. The strongest association was between Ca^2+^ decay time and contraction decay time (rho = 0.39, P = 0.067). Each point is one well, colored by condition and shaped by line. Panels **a**–**g** show the seven pairs tested, with Spearman ρ and P in each panel.

**Supplementary Fig. 6. Cell and nucleus size per condition and cell line.** Single-cell distributions of (**a**) cell area, (**b**) nucleus area in the calcium session (S1), (**c**) nucleus-to-cytoplasm area ratio and (**d**) nucleus area in the contraction session (S2), on a log10 axis. Brackets show Tukey-adjusted post hoc contrasts against control from the mixed models (* P < 0.05, ** P < 0.01, *** P < 0.001, ns, not significant). The cell line main effect of each model is given above each panel. Numbers above each condition give the number of cells. Cells outside the plotted range are not shown.

### Supplementary Tables

**Supplementary Table 1. Well-level quality control.**

For every imaged well, the number of segmented cells, the number and percentage of electrically responsive and irregular cells, the percentage of cells flagged as responders by CyteSeer (calcium session only), the QC status and the reason for exclusion. Wells were excluded if they contained fewer than 10 electrically responsive cells or, in the calcium session, if fewer than 71.0% of cells were CyteSeer responders (Otsu threshold). Wells that beat at twice the pacing frequency were excluded by hand. Abbreviations: QC, quality control.

**Supplementary Table 2. Cluster-number diagnostics.**

Within-cluster sum of squares and mean silhouette width for k-means solutions with k = 2 to 6, per imaging session, on z-scored clustering variables. Silhouette widths were computed on a random subsample of 5,000 cells. Both sessions were clustered with k = 3. Abbreviations: k, number of clusters.

**Supplementary Table 3. Linear mixed-model ANOVA.**

Type III F tests (Satterthwaite denominator degrees of freedom) for condition, cell line and their interaction in the model parameter ∼ condition × cell line + (1 | well), fitted on log10-transformed values of each parameter. Durations longer than the 1,000 ms pacing interval were treated as missing. Abbreviations: df, degrees of freedom.

**Supplementary Table 4. Post hoc pairwise comparisons.**

Pairwise comparisons between conditions within each cell line from the models in Supplementary Table 3 (estimated marginal means, Tukey adjustment, Kenward-Roger degrees of freedom). Estimates are differences on the log10 scale, and the ratio gives the first condition relative to the second. Abbreviations: SE, standard error; df, degrees of freedom.

**Supplementary Table 5. Functional subpopulations: profiles, composition and well-level tests.**

(A) Cluster centres in original units and cluster sizes. (B) Number and percentage of cells per cluster for each condition and cell line, the median percentage across wells and the change from control. (C) Kruskal-Wallis tests of the per-well cluster percentage across conditions, per cell line and cluster, with Dunn post hoc tests against control (Holm adjustment). Calcium clusters are numbered by kinetics (1 intermediate, 2 fastest, 3 slowest), contraction clusters by amplitude and BODIPY (1 highest amplitude, 3 highest BODIPY). Abbreviations: KW, Kruskal-Wallis.

**Supplementary Table 6. Condition effects within clusters.**

Condition effect tested separately within each cluster with parameter ∼ condition + cell line + (1 | well) on log10-transformed values (Kruskal-Wallis on well means where the mixed model could not be estimated). Means are estimated marginal means back-transformed to original units, averaged over the cell line, with Dunnett-adjusted contrasts against control. Because the parameters were also clustering variables, these tests are exploratory.

