## Supplementary Figure 1 for "CardioShift: Resolving cell-to-cell functional heterogeneity in hiPSC-derived cardiomyocytes from high-throughput single-cell imaging"

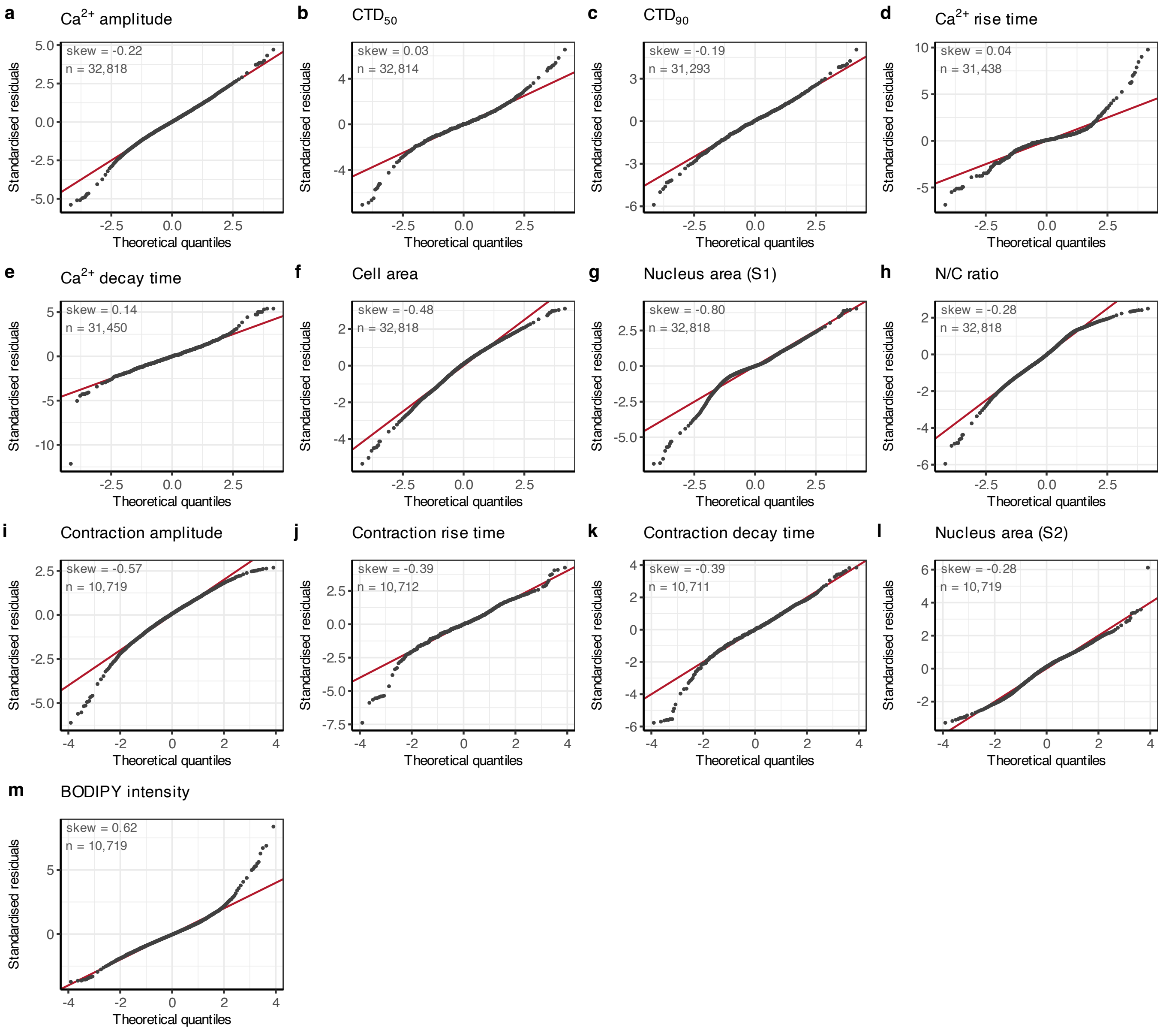
a Ca2+ amplitude
skew = -0.22
n = 32,818
b CTD50
skew = 0.03
n = 32,814
c CTD90
skew = -0.19
n = 31,293
d Ca2+ rise time
skew = 0.04
n = 31,438
e Ca2+ decay time
skew = 0.14
n = 31,450
f Cell area
skew = -0.48
n = 32,818
g Nucleus area (S1)
skew = -0.80
n = 32,818
h N/C ratio
skew = -0.28
n = 32,818
i Contraction amplitude
skew = -0.57
n = 10,719
j Contraction rise time
skew = -0.39
n = 10,712
k Contraction decay time
skew = -0.39
n = 10,711
l Nucleus area (S2)
skew = -0.28
n = 10,719
m BODIPY intensity
skew = 0.62
n = 10,719
Standardised residuals
Theoretical quantiles
