## Supplementary Figure 2 for "CardioShift: Resolving cell-to-cell functional heterogeneity in hiPSC-derived cardiomyocytes from high-throughput single-cell imaging"

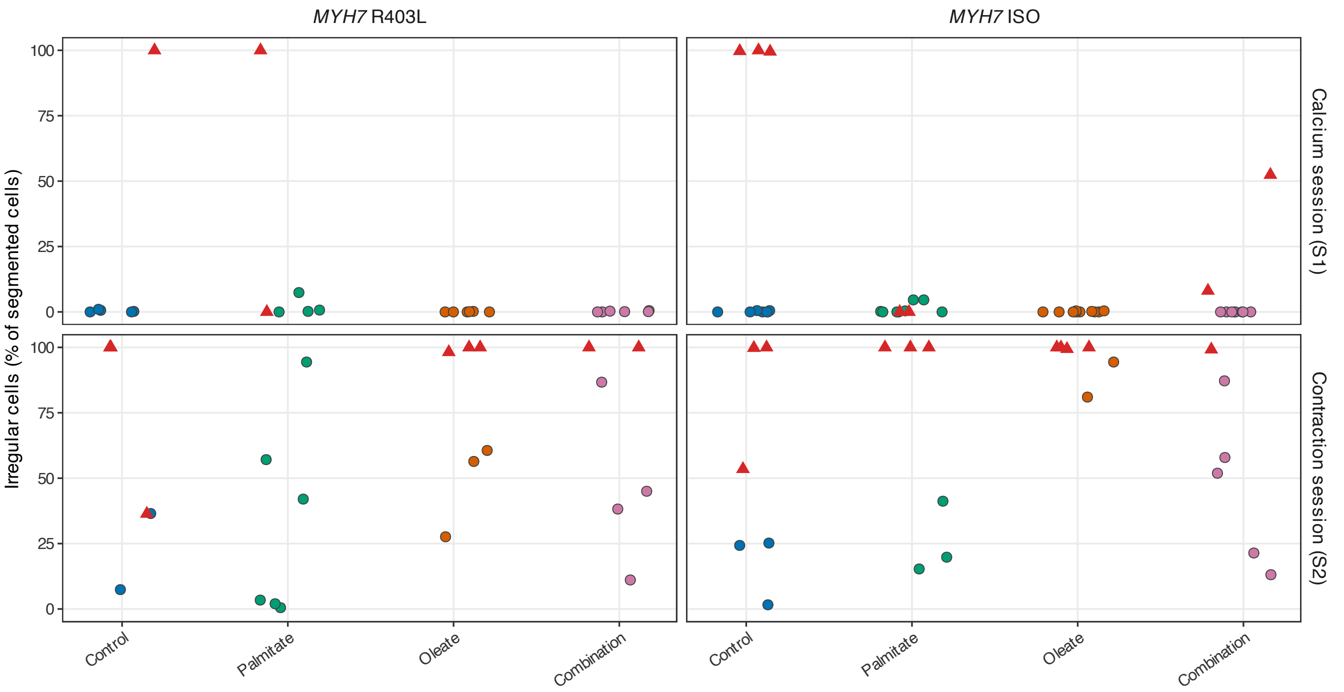
MYH7 R403L
MYH7 ISO
Irregular cells (% of segmented cells)
100
75
50
25
0
Calcium session (S1)
Contraction session (S2)
Control
Palmitate
Oleate
Combination
