## Supplementary figures and images for "CardioShift: Resolving cell-to-cell functional heterogeneity in hiPSC-derived cardiomyocytes from high-throughput single-cell imaging"

### Supplementary Figure 3

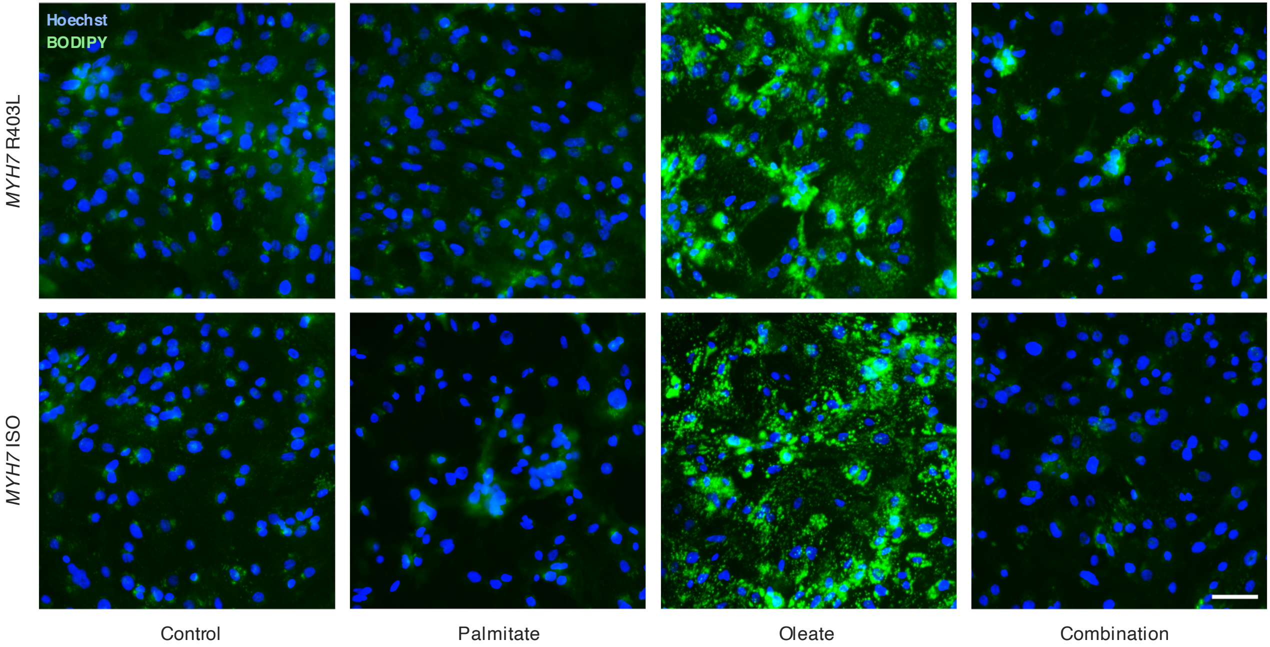
Hoechst
BODIPY
MYH7 R403L
MYH7 ISO
Control
Palmitate
Oleate
Combination
