## Supplementary Figure 4 for "CardioShift: Resolving cell-to-cell functional heterogeneity in hiPSC-derived cardiomyocytes from high-throughput single-cell imaging"

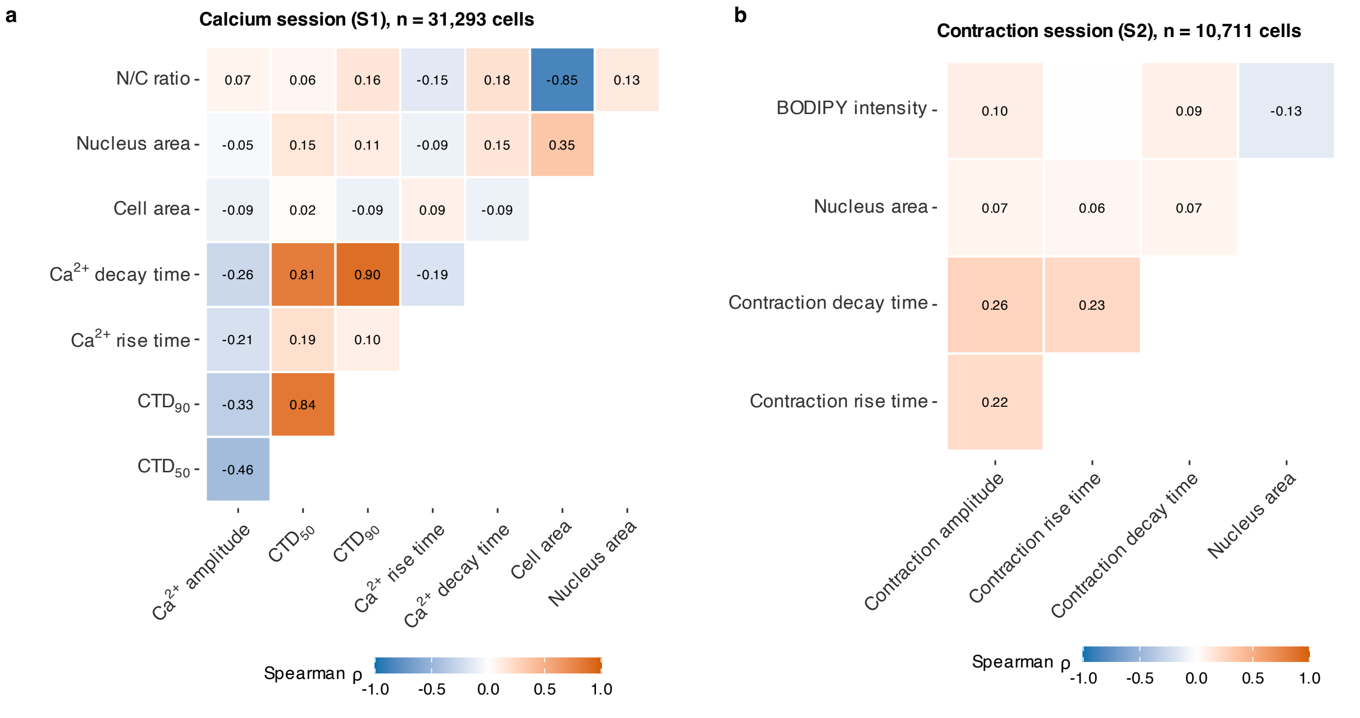
a
Calcium session (S1), n = 31,293 cells
N/C ratio
Nucleus area
Cell area
Ca2+ decay time
Ca2+ rise time
CTD90
CTD50
0.07
0.06
0.16
-0.15
0.18
-0.85
0.13
-0.05
0.15
0.11
-0.09
0.15
0.35
-0.09
0.02
-0.09
0.09
-0.09
-0.26
0.81
0.90
-0.19
-0.21
0.19
0.10
-0.33
0.84
-0.46
Ca2+ amplitude
CTD50
CTD90
Ca2+ rise time
Ca2+ decay time
Cell area
Nucleus area
Spearman ρ
-1.0
-0.5
0.0
0.5
1.0
b
Contraction session (S2), n = 10,711 cells
BODIPY intensity
Nucleus area
Contraction decay time
Contraction rise time
0.10
0.09
-0.13
0.07
0.06
0.07
0.26
0.23
0.22
Contraction amplitude
Contraction rise time
Contraction decay time
Nucleus area
Spearman ρ
-1.0
-0.5
0.0
0.5
1.0
