## Supplementary Figure 5 for "CardioShift: Resolving cell-to-cell functional heterogeneity in hiPSC-derived cardiomyocytes from high-throughput single-cell imaging"

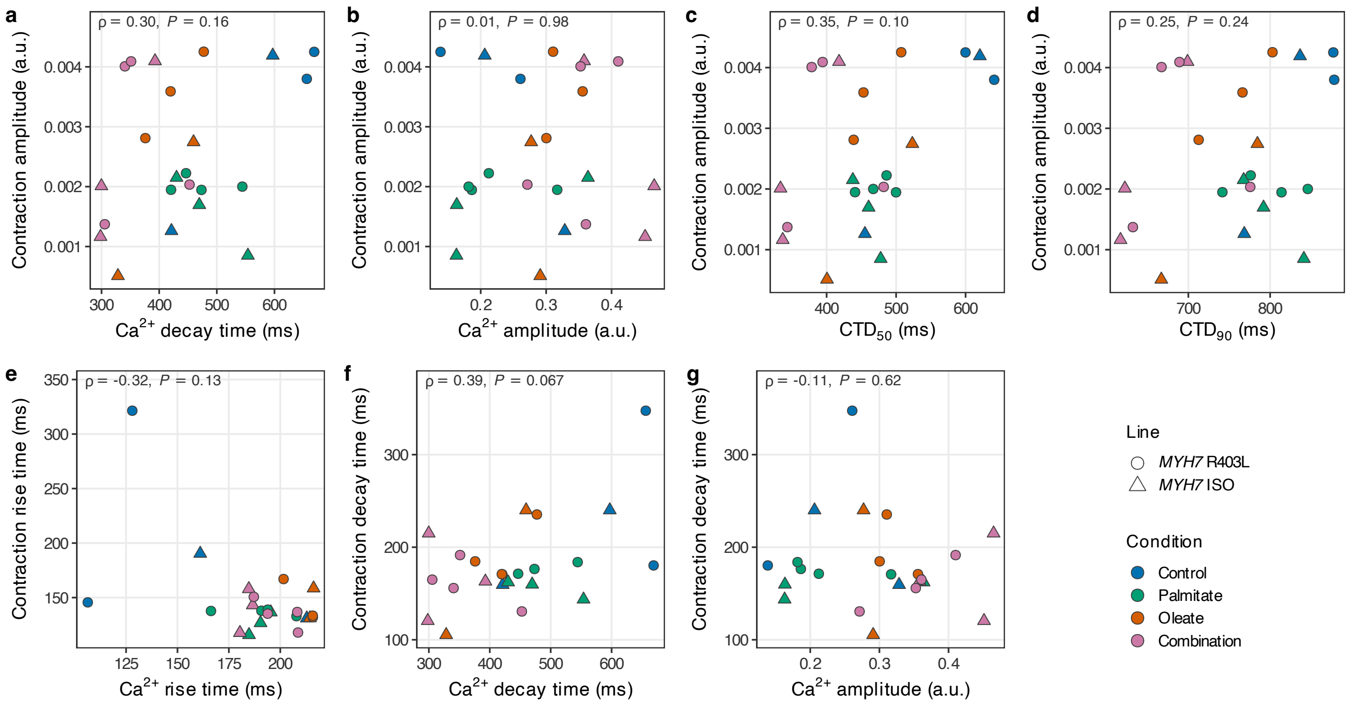

a
ρ = 0.30, P = 0.16
Contraction amplitude (a.u.)
Ca2+ decay time (ms)
b
ρ = 0.01, P = 0.98
Contraction amplitude (a.u.)
Ca2+ amplitude (a.u.)
c
ρ = 0.35, P = 0.10
Contraction amplitude (a.u.)
CTD50 (ms)
d
ρ = 0.25, P = 0.24
Contraction amplitude (a.u.)
CTD90 (ms)
e
ρ = -0.32, P = 0.13
Contraction rise time (ms)
Ca2+ rise time (ms)
f
ρ = 0.39, P = 0.067
Contraction decay time (ms)
Ca2+ decay time (ms)
g
ρ = -0.11, P = 0.62
Contraction decay time (ms)
Ca2+ amplitude (a.u.)
Line
MYH7 R403L
MYH7 ISO
Condition
Control
Palmitate
Oleate
Combination
