## Supplementary Figure 6 for "CardioShift: Resolving cell-to-cell functional heterogeneity in hiPSC-derived cardiomyocytes from high-throughput single-cell imaging"

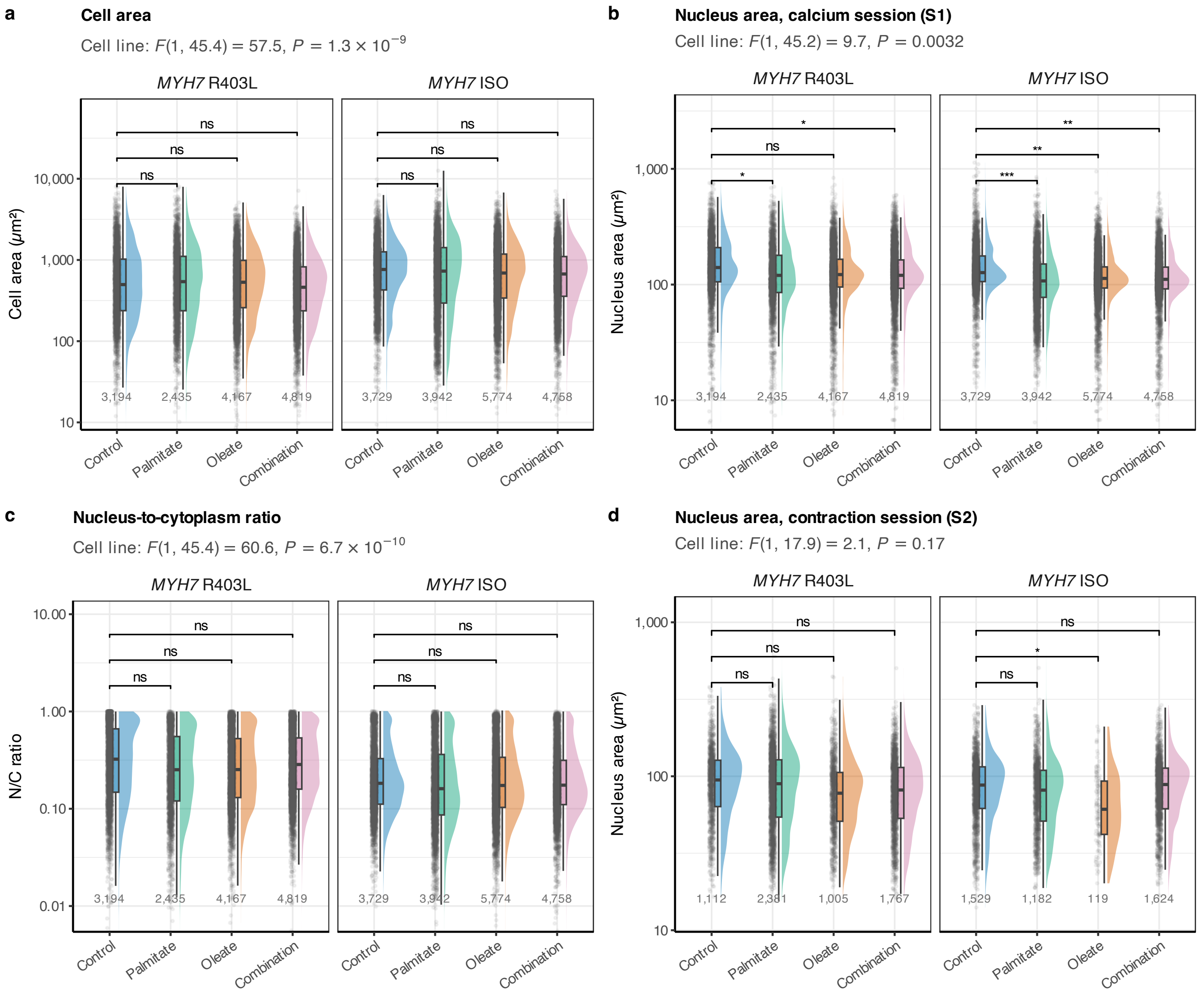

a
Cell area
Cell line: F(1, 45.4) = 57.5, P = 1.3 × 10−9
MYH7 R403L
MYH7 ISO
Cell area (μm²)
10,000
1,000
100
10
ns
ns
ns
3,194
2,435
4,167
4,819
3,729
3,942
5,774
4,758
Control
Palmitate
Oleate
Combination
b
Nucleus area, calcium session (S1)
Cell line: F(1, 45.2) = 9.7, P = 0.0032
Nucleus area (μm²)
1,000
100
10
*
**
***
c
Nucleus-to-cytoplasm ratio
Cell line: F(1, 45.4) = 60.6, P = 6.7 × 10−10
N/C ratio
10.00
1.00
0.10
0.01
d
Nucleus area, contraction session (S2)
Cell line: F(1, 17.9) = 2.1, P = 0.17
1,112
2,381
1,005
1,767
1,529
1,182
119
1,624
